# Structures of the Cyanobacterial Phycobilisome

**DOI:** 10.1101/2021.11.15.468712

**Authors:** Paul V. Sauer, Maria Agustina Dominguez-Martin, Henning Kirst, Markus Sutter, David Bina, Basil J. Greber, Eva Nogales, Tomáš Polívka, Cheryl A. Kerfeld

**Affiliations:** QB3 Institute, University of California, Berkeley, CA, USA; Howard Hughes Medical Institute, University of California, Berkeley, CA, USA; Environmental Genomics and Systems Biology Division, Lawrence Berkeley National Laboratory, Berkeley, CA 94720, USA.; Molecular Biophysics and Integrated Bioimaging Division, Lawrence Berkeley National Laboratory, Berkeley, CA 94720, USA.; MSU-DOE Plant Research Laboratory, Michigan State University, East Lansing, MI 48824, USA.; Faculty of Science, University of South Bohemia, Ceske Budejovice, Czech Republic; Biology Centre of the Czech Academy of Sciences, Ceske Budejovice, Czech Republic; Division of Structural Biology, Institute of Cancer Research, London, UK; Department of Molecular and Cellular Biology, University of California, Berkeley, CA, USA.; Department of Biochemistry and Molecular Biology, Michigan State University, East Lansing, MI 48824, USA

## Abstract

The phycobilisome is an elaborate antenna that is responsible for light-harvesting in cyanobacteria and red-algae. This large macromolecular complex captures incident sunlight and transfers the energy via a network of pigment molecules called bilins to the photosynthetic reaction centers. The phycobilisome of the model organism Synechocystis PCC 6803 consists of a core to which six rods are attached but its detailed molecular architecture and regulation in response to environmental conditions is not well understood. Here we present cryo-electron microscopy structures of the 6.2 MDa phycobilisome from Synechocystis PCC 6803 resolved at 2.1 Å (rods) to 2.7 Å (core), revealing three distinct conformations, two previously unknown. We found that two of the rods are mobile and can switch conformation within the complex, revealing a layer of regulation not described previously. In addition, we found a novel linker protein in the structure, that may represent a long-sought subunit that tethers the phycobilisome to the thylakoid membrane. Finally, we show how excitation energy is transferred within the phycobilisome and correlate our structures with known spectroscopic properties. Together, our results provide detailed insights into the biophysical underpinnings of cyanobacterial light harvesting and lay the foundation for bioengineering of future phycobilisome variants and artificial light harvesting systems.

---

Cyanobacteria are the most abundant and ecophysiologically diverse primary producers on Earth. Oxygenic photosynthesis is an ancient cyanobacterial innovation that allowed complex life to emerge^1^, and today these organisms are being established as platforms for green biotechnologies^2^. Cyanobacterial phycobilisomes (PBSs)^3–5^ are massive pigment-protein complexes that can constitute up to half of the soluble protein content of the cell^6^. The two major PBS substructures, the core and the rod, are composed of phycobiliproteins, and colorless linker proteins (LPs)^7^. Phycobilin pigments are covalently bound to the phycobiliproteins^8–14^, which assemble into disc-like trimers (αβ)3 or hexamers (αβ)6. These discs are chained together in the rods and core cylinders by LPs and organized into a complete PBS^15–17^. The association of pigments with protein tunes the pigments’ energetic properties to establish an energy cascade to the photosynthetic reaction centers that is both extremely fast and highly efficient (ca. 95%)^18, 19^. Four morphological types of PBS are known: hemiellipsoidal^16, 20, 21^, block-type^15^, hemidiscoidal^10, 22–25^ and bundle-type^26^. *Synechocystis* PCC 6803 is the primary model system for molecular and spectroscopic studies ^27^ containing the most widespread type of PBS, the hemidiscoidal^8, 22, 25, 28, 29^, which is also the evolutionary antecedent of all types of PBS^15^. We report three cryo-electron microscopy (cryo-EM) structures of the 6.2 MDa PBS from *Synechocystis* PCC 6803, representing distinct conformers of rod arrangements: the canonical ‘up-up’ (2.7 Å), and two previously undescribed, ‘up-down’ (2.8 Å), and ‘down-down’ (3.5 Å) conformations. We also describe the structure of all the linkers, one of them previously unknown, the arrangement of all bilins, and elucidate the likely energy transfer pathway.

## Structural overview

Purified and intact PBS complex was biotinylated and subjected to cryo-EM using streptavidin affinity grids^30^ to reduce preferential particle orientations and to increase stability (Methods, Extended Data Fig. 1, Extended Data Table 1, 2). We determined the structure of three distinct PBS conformations at resolutions ranging from 2.7 to 3.5 Å for the entire complexes, enabling us to build all pigments and protein chains (Fig. 1, Extended Data Fig. 2, 3, and Extended Data Table 3). As previously reported^22^, the rods are flexible in nature and therefore appear smeared in the reconstructions. Therefore, the rods were processed and reconstructed independently, yielding a resolution of 2.1 Å. The rods cannot be further sorted into distinct classes, suggesting that all rods within one complex are identical (Extended Data Fig. 2c).

**Figure 1:**
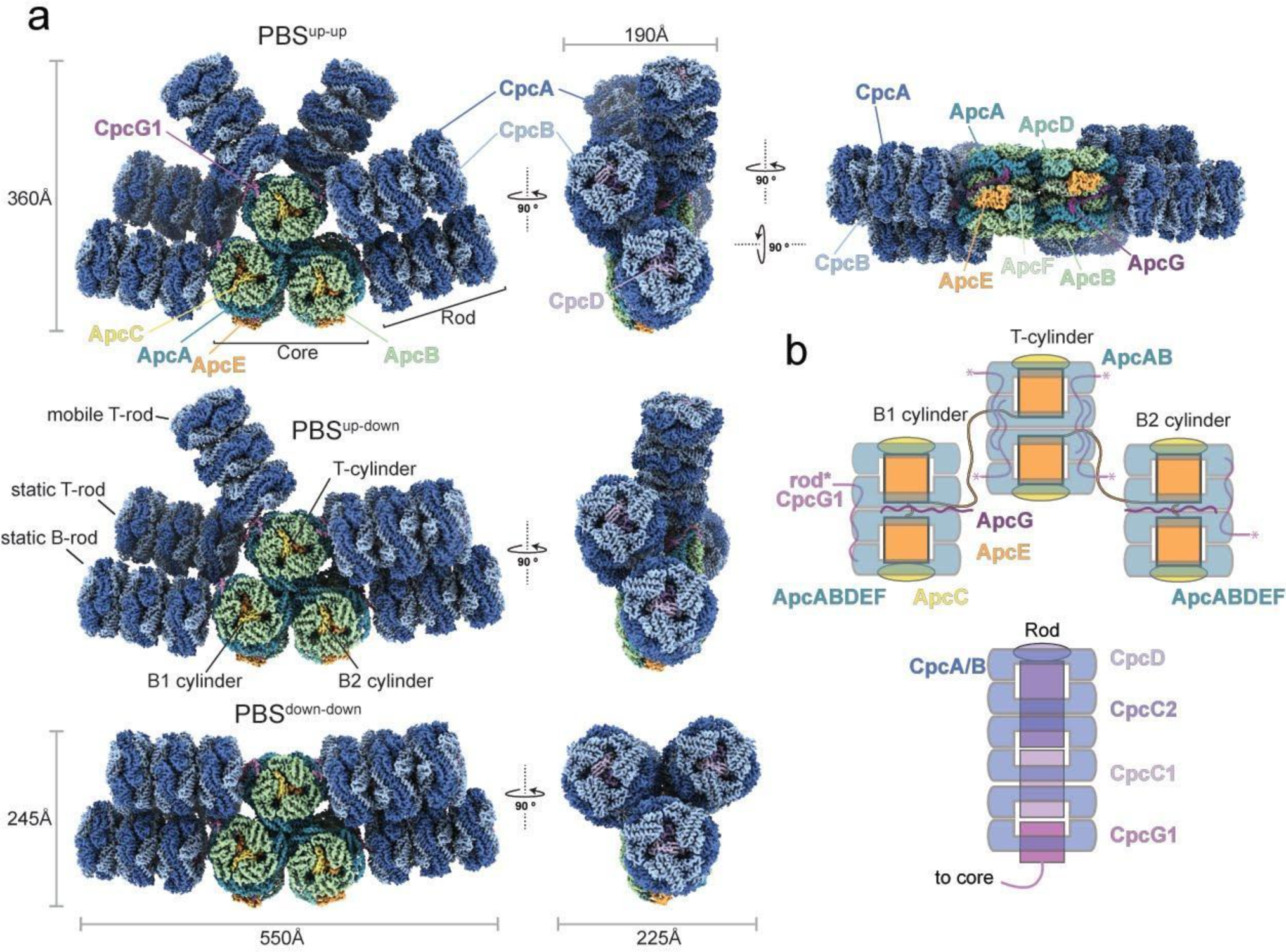
Structural characterization of the phycobilisome from Synechocystis PCC 6803. **a**, Composite cryo-EM density maps representing the three PBS conformations: Up-up (top), up-down (middle), down-down (bottom). Individual maps and resolution for the three conformations and the rod can be found in Extended Data Fig. 2. **b**, Schematic overview of PBS connectivity and subunit arrangement of the core (top) and the rods (bottom).

**Figure 2:**
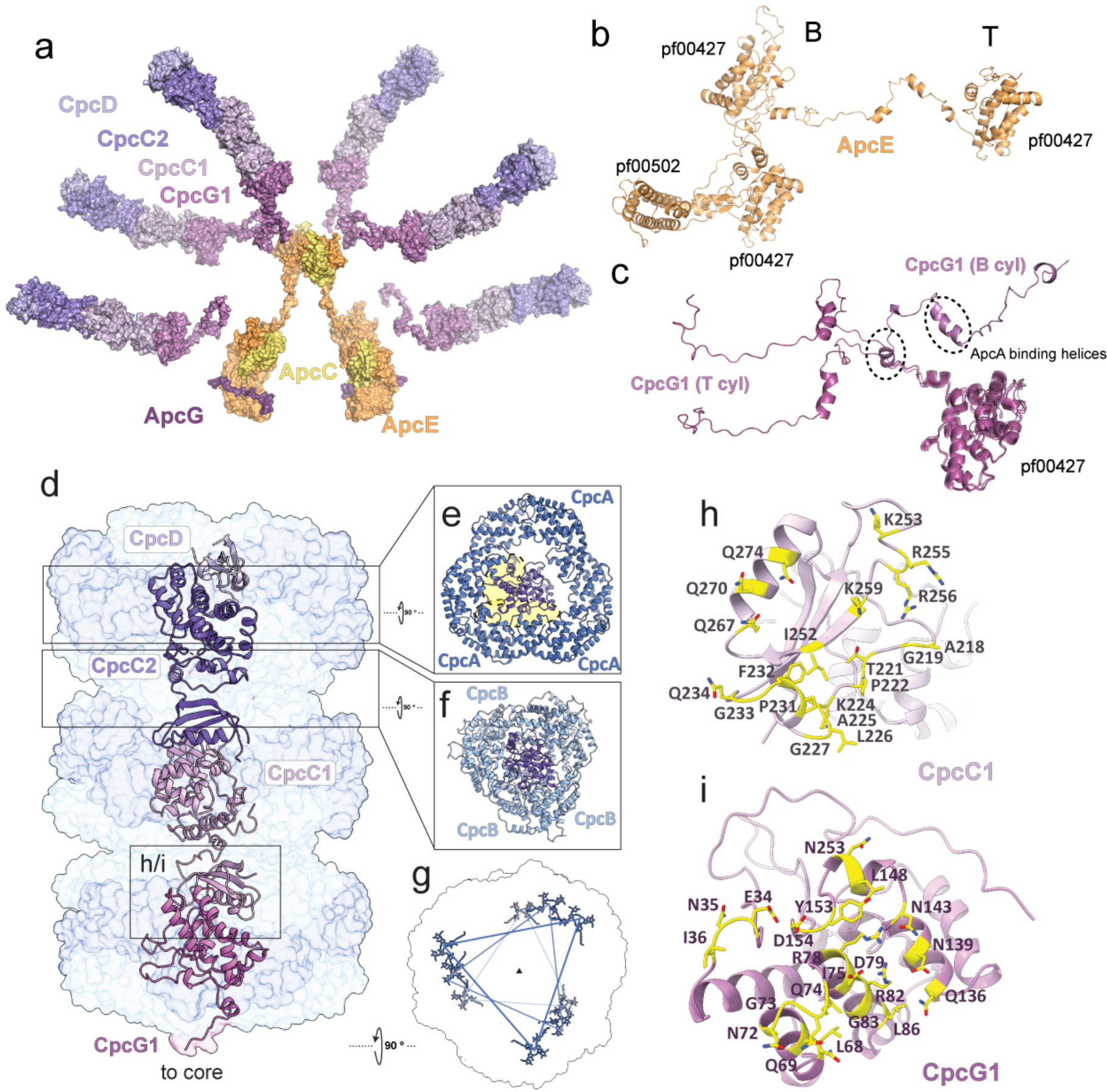
Core and rod linkers. **a**, Surface representation of the linker skeleton of the entire PBS. **b**, ApcE with its constituent domains pf00502 and pf00427. **c**, CpcG1 linker conformations from top (T) and bottom (B) cylinders in the up up conformation. **d**, Linker proteins within a rod. CpcC1/2 contain a pf00427 and pf01383 domain, CpcD only a pf01383 domain **e**, Interface of CpcC2 with CpcA proteins. Predominant contacts between the linker and CpcA proteins are highlighted in yellow. **f**, CpcC1/2 interact more closely with CpcB than CpcA. **g**, Bilin arrangement within the rod showing spiraling pattern. **h** and **i**, CpcC1 and CpcG1 amino acids interfaces along the long axis of the rod highlighted in yellow.

**Figure 3:**
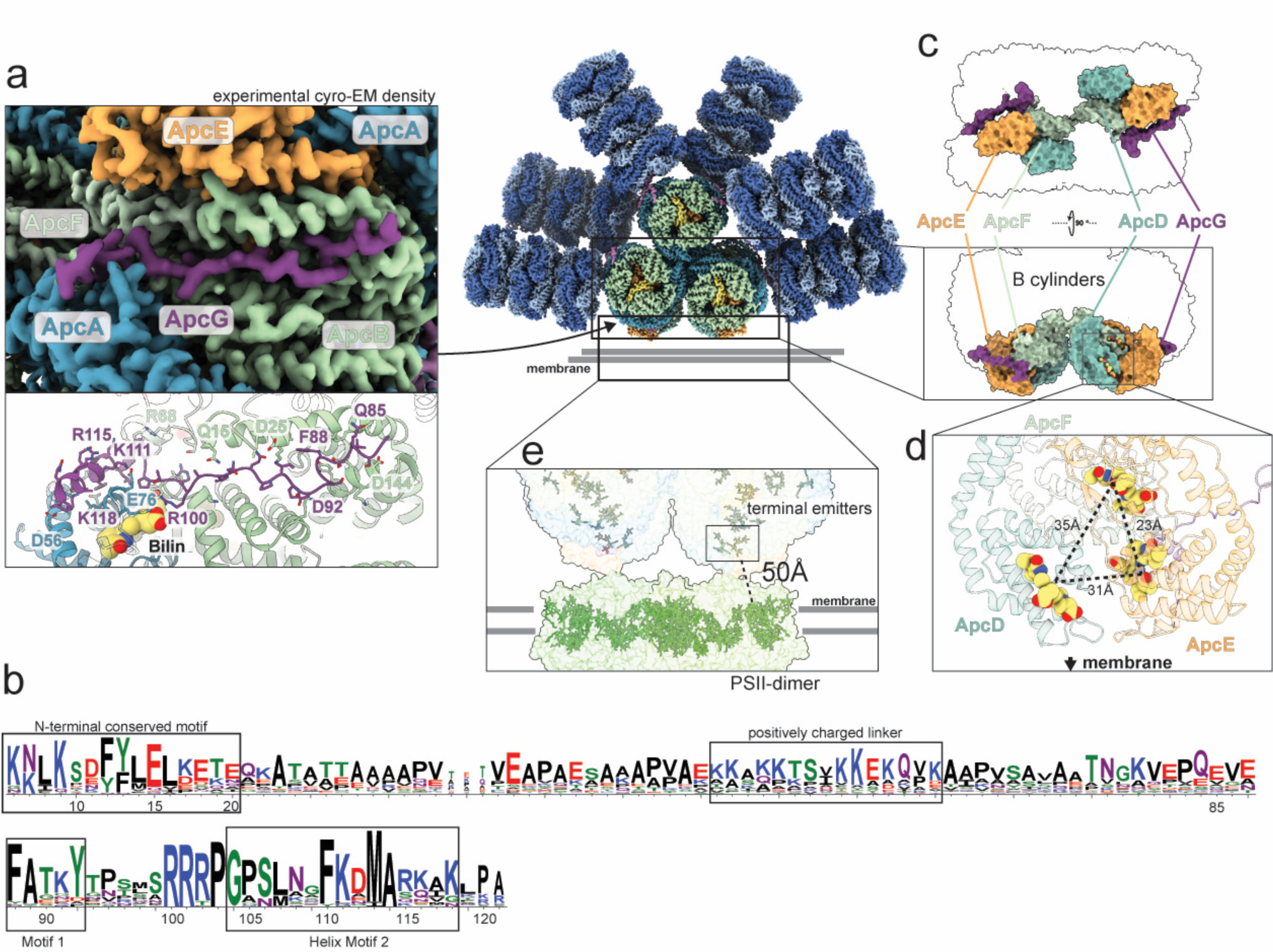
A newly identified linker ApcG and Terminal emitters. **a**, Position and interaction of ApcG within the PBS. **b**, Sequence conservation logo of ApcG homologs. The numbers refer to residue positions in Synechocystis PCC 6803 ApcG. ApcG contains three conserved sequence motifs including a short helix that binds in a central groove of ApcA like CpcG1. **c**, Position and orientation of the subunits ApcD, ApcE, ApcF and ApcG within the core of the PBS. Only the phycobiliprotein domain of ApcE is rendered. **d**, Orientation and distances of the PCBs of ApcD, ApcE and ApcF. **e**, Rigid body modelling of our PBS core structure onto PSII, using EMDB 2822 as a reference.

The PBS core consists of three cylinders, a top cylinder (T) stacked on top of two basal cylinders (B1, B2) (Fig. 1, Extended Data Fig. 4). Each of the two B cylinders have one rod attached while four rods are attached to the T cylinder. The core consists of 80 polypeptide chains and 72 phycocyanobilins (PCB), and each rod contains 40 protein subunits and 54 PCBs, adding up to a total of 320 proteins and 396 bilin molecules that amount to a molecular weight of 6.2 MDa. Extended Data Figure 4 and Extended Data Table 1 provide an inventory of the subunits, their component protein family (pfam) domains and functional characteristics.

**Figure 4:**
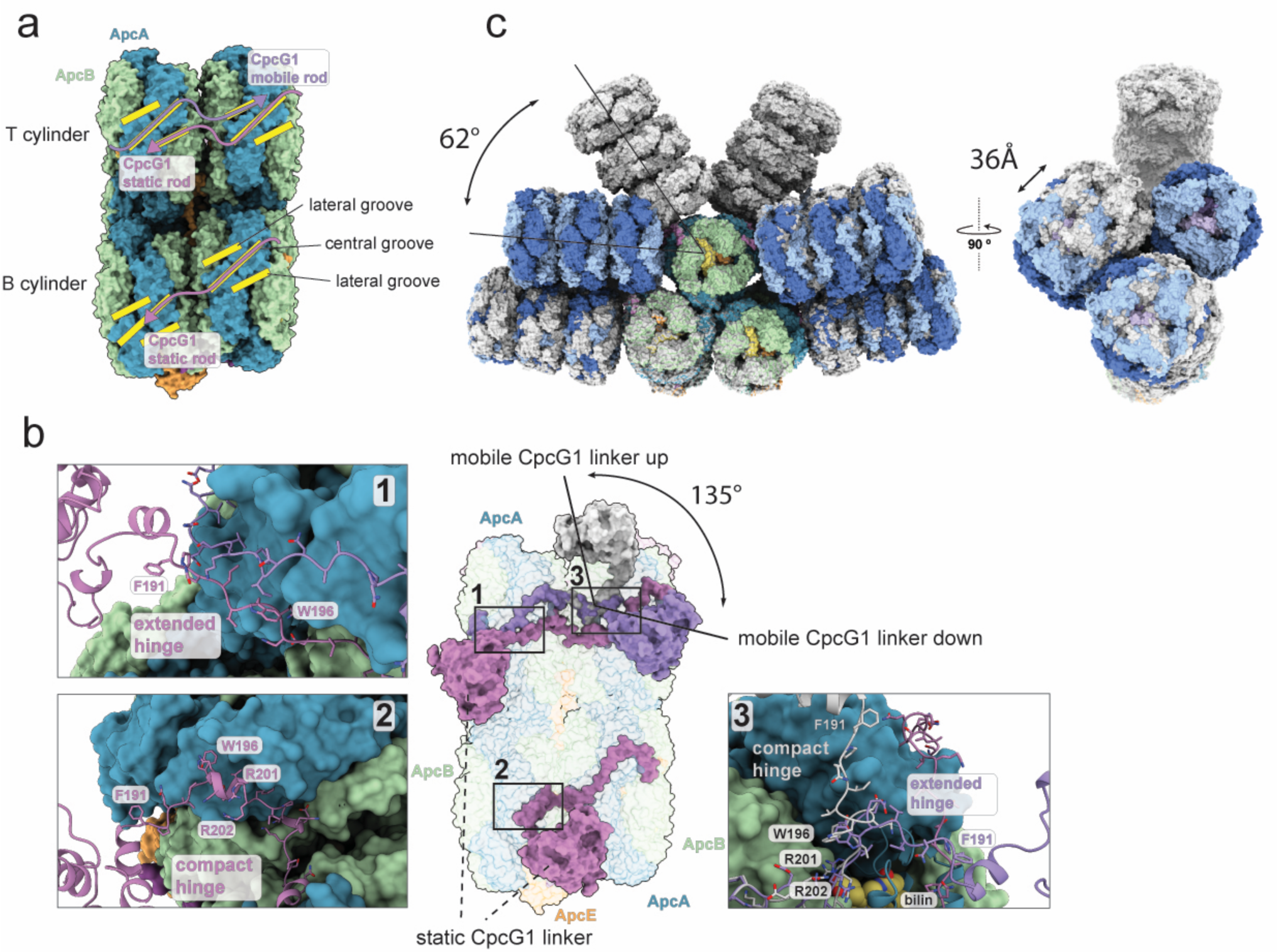
Structural basis for conformational switching. **a** Side view of the PBS core omitting the rods showing the groove network that organizes CpcG1 linker attachment. **b**, Same side view of the PBS core emphasizing the CpcG1 position. Center panel shows an overview while numbered insets show the different hinge conformations. Near the hinge of the mobile rod is a PCB molecule that is contacted by R201 of CpcG1. **c**, Superimposition of PBSdown-down (in color) and PBSupup (light grey) showing the large conformational differences between the two.

In contrast to earlier studies, our structures include three different, major rod conformations; two have not been visualized before. They are distinguished by alternate positioning of two of the four T rods (hence referred to as mobile T rods, as opposed to the static T rods). In the ‘up-up’ conformation, both mobile rods are tilted upwards, corresponding to the canonical hemidiscoidal PBS. Two previously unreported conformers were also present: the ‘up-down’ conformation, where one of the mobile rods moved downwards to assume a position parallel to the other rods and the ‘down-down’ conformation, in which all the T cylinder rods are in a down position, reducing the ‘height’ of the complex by more than 100 Å and giving the PBS a more compact profile (Fig. 1a).

## Linker architecture and identification of a new linker component, ApcG

Both the core and the rods contain linker proteins in their central cavities that provide scaffolding and organize the arrangement of the Apc and Cpc subunits, thereby ensuring the proper orientation and spacing of the pigments (Fig. 1b, 2a). Central to the organization are two copies of ApcE (Fig. 2a, b, Extended Data Fig. 5a) which form the major scaffold for the core by connecting the bottom with the top cylinders^15–17^. The PB loop (residues 87-129) is disordered and could not be modeled, but all other ApcE domains are well resolved. The three cylinders are ‘capped’ by six copies of ApcC that interact with ApcE (Fig. 2a, Extended Data Fig. 5b).

**Figure 5:**
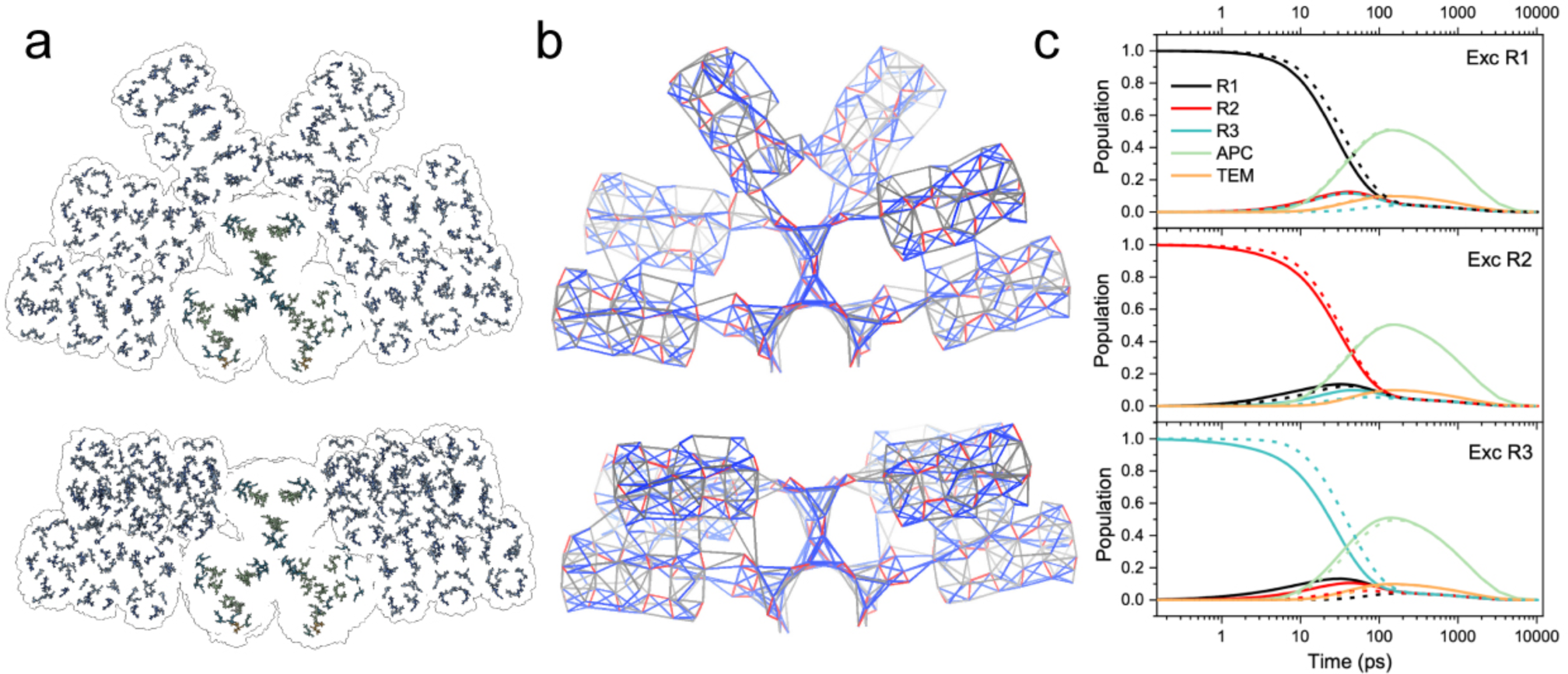
Energy flow through the PBS. **a**, Pigment distribution within PBSup-up (top) and PBSdown-down(bottom). **b**, Maps of individual inter-pigment energy transfer rates of PBS^up-up^ (top) and PBS^down-down^ (bottom). The color bars correspond to rates faster than 1 ps (red), in the 1-10 ps range (blue) and in 10-20 ps range (grey). Inter-pigment rates slower than 20 ps are omitted. **c**, Energy flow through down-down (solid) and up-up (dashed) PBS configurations. The excitation is placed either on the mobile rod R1 (top), middle rod R2 (middle) or bottom rod R3 (bottom). The curves monitor the evolution of excited state population of pigments located in rod, core (APC), and terminal emitters (TEM).

In the rods, there are four linker proteins (Fig. 2 a, d). CpcG1 is located at the core-proximal side. Its 60 residue C-terminal extension attaches it to the PBS core (Figure 2 c, d, Extended Data Fig. 5c). The homologues CpcC1 and CpcC2 form the bulk of the interior of the rods (Fig. 2d, Extended Data Fig. 5 d). The interdomain linker of CpcC1 is 15 residues longer than in CpcC2, allowing for a 70° rotation of the pf01383 domain of CpcC1 compared to CpcC2 (Extended Data Fig. 5d). This results in an apparent 10° rotation between each CpcA/B hexamer (Fig. 2g). Even though all linker proteins within the rod employ the same two domains to connect to one another, differences in the binding interface of each domain pair define a unique architecture for the correct assembly of the entire rod (Fig. 2 h, i, Extended Data Fig. 5 e-g).

A previously unknown linker protein (*sll1873*) was discovered in the cryo-EM density and identified by mass spectrometry (Fig. 3a, Extended Data Table 2). We denote it ApcG or LC10 in continuation of the established nomenclature. It is found only on the bottom two cylinders extending toward the membrane-facing side. An HMM search reveals that homologues are found in about 80% of cyanobacterial genomes. ApcG contains two conserved sequence motifs including a short helix that binds in a groove on ApcA like CpcG1 (Fig. 3b, Extended Data Fig. 5b). The highly conserved residue R100 of ApcG directly contacts the PCB of ApcA in the bottom cylinder, suggesting that it may finetune its spectroscopic properties. It has previously been suggested that the PBS interacts with lipid head groups^31^; ApcG could mediate this interaction through its positively charged linker region (Fig. 3b). Given its location close to the terminal emitters and the membrane, we propose that ApcG likely plays a key role in PBS localization and energy transfer to PSII.

## Terminal emitters

At the membrane facing side of the PBS, the terminal chromophores of ApcD and ApcE are responsible for transferring the absorbed energy to the reaction centers (Fig. 3c, Extended Data Fig. 4). ApcF has been shown to play a crucial role in energy migration to ApcE and also to impact state transitions^32, 33^. The center-to-center distance between the PCBs of ApcE and ApcF is 23 Å, and 35 Å between the PCBs of ApcD and ApcF (Fig. 3d). While these distances fall well within the range of other PCB distances, their local environments are likely responsible for the observed shift in fluorescence emission of ApcE and ApcF to 680 nm (as opposed to 660 nm for ApcA). Rigid body docking of our structure into a negative stain map of the related *Anabaena sp.* PBS bound to PSII^25^ suggests that the closest inter-pigment distance between the PBS and PSII might be as large as ∼50 Å (Fig. 3e). The requirement to spectroscopically bridge such a distance might be one of the drivers for the diversification of ApcD, E and F around the terminal chromophores of the PBS and leading to their unique properties.

## Core-rod linker architecture and conformational switching

Rod attachment is mediated by CpcG1, which protrudes from the central cavity of each rod and latches onto the surface of the core cylinders. The PBS core accommodates the CpcG1 linkers through a network of grooves that span the surfaces of ApcA and ApcB and which can be divided into central and lateral grooves (Fig. 4a, Extended Data Figure 6a). Unlike the algal PBS complexes, grooves from separate discs can align to create a curving binding path for the CpcG1 linkers (Figure 4a).

The conformations of the CpcG1 linkers correlate within the three rod (Fig. 4b). When comparing the linkers of the top and bottom static rods it is apparent that the linker can adopt one of two different conformations: extended or compact. In the compact state, a conserved six amino acid region (W196-R201), which we define as the ‘hinge’ forms a small coil or helix that changes the local geometry of the linker relative to the extended state (Fig. 2c, 4b). Of the two static rods, the hinge of the upper rod adopts the extended conformation, while the hinge of the lower rod is in the compact conformation (Fig 4b). As a consequence of this conformational difference, the globular domain of the lower CpcG1 is rotated with respect to the globular domain of the upper CpcG1. This rotation is locked in place only for the static rods, the mobile rod however can toggle between these two states which are determined by the conformation of the hinge. Superposing the position of CpcG1 of the mobile rod in its ‘down’ conformation with its own ‘up’ conformation reveals that the movement of the mobile rod between ‘down’ and ‘up’ corresponds to the conformational change of the linker between compact and extended. As a result, the globular CpcG1 domain along with the entire rod rotates by 135° around its long axis when swinging from ‘down’ to ‘up’ (Fig. 4b). Most importantly, this switch is accommodated by a 62° upwards swing of the entire rod, which is the most striking difference between the ‘down’ and ‘up’ conformations (Fig. 4c). A similar conformational change from the mobile rod to the static rods is precluded because it would result in a clash of the static rods. The bottom static rod would swing upwards while the upper static rod would swing downwards. This mutual blockade alone is sufficient to explain the static nature of these two rods.

The mobile rod is held in place in its ‘down’ position through interactions with its static rod neighbor. Two Cpc(αβ)6 hexamers proximal to the PBS core interact through a salt bridge that is formed between R150 of one of the CpcB’s in the mobile rod and E131 of one of the CpcA’s in the static rod (Extended Data Fig. 6b).

The superposition of ‘down’ and ‘up’ conformations also reveals that the upper static rod experiences a 36 Å lateral displacement when the mobile rod is down but relaxes into its central position once the mobile rod swings upwards (Fig. 4c). Once in the central position, the static rod obstructs the downward movement of the mobile rod and stabilizes this conformation. Hence, in every case the mobile rod needs to overcome an energy barrier to move either direction.

## Energy transfer pathways in the different PBS conformations

To better understand the impact of the different conformations we computationally modeled the energy transfer within the phycobilisome based on our structural data. We assume energy transfer in the limit of Förster theory using a dipole-dipole approximation for inter-pigment couplings. The map of energy transfer rates for the PBS in the up-up conformation is shown in Fig. 5, demonstrating a robust system, in which excitation of essentially any pigment directs the excitation quickly to the terminal emitters. It also shows that the bottleneck is the rod-to-core energy transfer, as suggested by global fitting models of time-resolved fluorescence data^34^. Energy transfer between the pigments in the core, both within and between the cylinders, is more efficient than rod-to-core transfer, contradicting the global fitting models that predict slow inter-cylinder transfer^35^, but the overall flow of energy through the whole PBS obtained from our model matches the experimental data well (Extended Data Fig. 7a, b).

The individual rate constants shown in the map are used to calculate how long the excitation resides in each structural (e.g., rod or core) or spectroscopic unit (Methods, Extended Data Fig. 7a, b). Using this approach, we also investigated the energy transfer through both the up-up and the down-down PBS conformations by placing the excitation to the most distant disc of the respective rods (Fig. 5c). For both conformations, the excitation reaches the APC core in less than 100 ps, in agreement with experimental data (see Extended Data Fig. 7a, b). Moving the mobile rod to the down position (solid lines in Fig. 5c) facilitates inter-rod energy transfer, as the more packed conformation increases the probability of energy transfer between bilins from different rods. Excitation of the mobile rod in down-down conformation leads to about 10 % population of the lowest rod, while for the up-up conformation the population of this rod is essentially zero (Fig. 5c). The same effect occurs for excitation of the bottom rod; essentially no inter-rod energy transfer is observed for the up-up conformation, while a non-negligible population is found at the other rods in the down-down PBS (Fig. 5c). Thus, the more compact down-down conformation allows for more pathways to reach the core, slightly increasing the overall trapping efficiency. On the other hand, the fact that significant movement of the mobile rod keeps the overall flow of energy through the system virtually unchanged points to robustness of PBS with respect to light-harvesting capacity. The robustness of PBS is further underscored by comparison with the red algal PBS, which shows that the overall dynamics are comparable^15, 16, 36^.

We note that two PBS configurations differing by their fluorescence spectra have been identified by single molecule spectroscopy^37^. It is thus tempting to associate the up-up and down-down conformations with those two species. Here, however, the structures of up-up and down-down conformations do not yet reveal any influence on the structure of the PBS core that is needed to change the fluorescence of terminal emitters. Thus, the relation of the two species identified by different methods remains to be defined.

## Importance of conformational switching for light harvesting

The quality of our structural data and the various proportions of the three PBS conformations suggests that we have captured previously unknown conformations of the cyanobacterial antenna. Why have the different rod conformations not been described before? In retrospect, it becomes clear that during earlier attempts to visualize the cyanobacterial PBS different conformation were probably confounded with general flexibility^22^ or likely disregarded as damaged particles^38^. Other methods to determine the structural arrangement of the rods like crosslink-MS are not suited to distinguish between inter and intra-rod contacts and therefore are likewise blind to the conformations we identified^29, 39^. Because the static rods already display two different hinge conformations, it is unlikely that the rod movement is an artifact from sample preparation or imaging. The observed movement is directional and quantifiable, hallmarks of true biologically relevant conformational changes. It is tempting to assume that the mechanistic cause for the rod movement is coupled to the transfer of excitation energy through the PBS. Pigment molecules can be found near several residues critical for rod movement (Fig. 3d, Extended Fig. 6b), but further studies are now needed to explore whether any of the pigments within the PBS can indeed trigger a conformational change and dictate rod position. The newly described PBS architectures suggests a regulation of light harvesting at the level of supramolecular organization in the thylakoid membrane that is somehow regulated by rod movement.

We propose two scenarios for placing our structures in that context. PBS arrays efficiently pack light harvesting pigments on the surface of the thylakoid membrane and thus increase the overall photon absorption cross-section of a cell. Additionally, they create redundancy of light harvesting and photochemical energy conversion at the reaction centers and can thus alleviate energetic losses due to damaged PSII. Using the published structure of an array as a model^40^, we found that only the ‘up-up’ state of the PBS could be fitted into the array (Extended Data Fig. 7c). Modeling of the energetics of our structure in arrays indicates that lateral energy transfer between PBSs within an array is possible, mainly between adjacent PBS cores.

Under certain conditions, PBS arrays might need to be dismantled to rebalance PSII versus PSI excitation pressure. Given that the down-down and down-up PBS conformations cannot be fitted into arrays, (Extended Data Fig. 7d, e), a conformational change in one PBS unit could therefore help to break up such arrays by acting as a terminator or interrupter of PBS array formation.

The second scenario in which PBS conformation may regulate the balance between light harvesting and photoprotection is coupled with the activity of the photoreceptor OCP. OCP quenches the excitation energy by converting it to vibrational energy. Activated OCP binds to the lateral side of the PBS core in the space between the rods^41^. However, the binding site is completely blocked when the mobile rod is in the ‘down’ position. Therefore, the mobile rod controls access of OCP to the PBS core and determines whether the PBS can be quenched.

Having more than one mechanism could be a means for the cell to fine tune its light harvesting efficiency to adjust to altered metabolic and cellular requirements. Moreover, these scenarios are not mutually exclusive and could work concurrently, but on different time scales, to adjust light-harvesting in response to environmental changes. The OCP binding site is still accessible in PBS arrays^41^, and thus PBS arrays are readily available for OCP induced excitation energy quenching upon rapid changes in the environment. Supramolecular rearrangements of the photosynthetic machinery can then take place to rebalance light harvesting, which might involve rod movement and array dispersion. Several studies have revealed that excitation energy harvested at the PBS can be transferred to PSI^39^. Given that PSI does not require excitation energy quenching at the light-harvesting system, it can be hypothesized that the down-down position obscuring the OCP binding site might be involved mainly in transferring its harvested energy to PSI. In fact, in such a scenario, OCP induced quenching would be counterproductive.

Here, we present the first high-resolution structures of cyanobacterial phycobilisomes, revealing an unexpected variety of conformational states that help to collect and deliver light energy to photosynthetic reaction centers. Further analysis of these structures will inform future experiments that will help us to understand and harness the power of these large light harvesting machines.

## Methods

### Preparation of the *Synechocystis* PCC 6803 Phycobilisome

#### Synechocystis

PCC 6803 were grown photoautotrophically in a BG11 medium. Cells were kept in a rotary shaker (100 rpm) at 30°C, under 3% CO2 enrichment, illuminated by white fluorescent lamps with a total intensity of about 30 μmol photons m^-2^ s^-1^. The protocol used for PBS isolation was based on^42, 43^. The cells were collected at 8,000 rpm, 10 min at room temperature. The pellet was resuspended in 10 mL of PBS isolation buffer (0.75 M potassium phosphate buffer pH 7.5, 1 mM EDTA, 0.5 mM PMSF) per 2 g of fresh weight of cells. Then, the cells were washed twice by centrifugation at 30,000 *g* during 15 min. The cells were broken through French Press at 1,000 psi three times. The supernatant was incubated in the presence of 2% (v/v) Triton X-100 under dim stirring, at 23°C, 15-20 min. The cell debris and the aggregates were removed by ultracentrifugation at 20,000 rpm (Ti-70 rotor) for 20 min at room temperature. The dark blue supernatant was directly loaded onto a discontinuous sucrose gradient (0.5, 0.75, 1, and 1.5 M sucrose layers in 0.75 M potassium phosphate buffer, pH 7.5), and spun at 150,000 *g*, 23°C, for 16 h using a SW28 rotor. The intact PBS was recovered from the 0.75 – 1 M interface of the sucrose gradient. The PBS were buffer exchanged into 0.75 M potassium phosphate buffer pH 7.5 using Amicon (30 kDa). The samples were kept at room temperature for further use.

### Protein separation

The isolated PBS samples were concentrated by precipitation with 20% (v/v) trichloroacetic acid prior to loading on a sodium dodecyl sulfate-polyacrylamide gel electrophoresis (SDS-PAGE) 5-20% gradient gel. The gels were run for 25 min at 240 V and were stained by Coomassie Brilliant Blue. The protein composition of the SDS-PAGE confirms the presence of all PBS subunits (Extended Data Tables 1, 2).

### Absorption and fluorescence spectrum measurement

PBS ultraviolet–visible (UV-vis) absorption spectra were collected with a Cary 60 spectrophotometer (Agilent). The fluorescence emission spectra of the PBS were recorded at room temperature from 600 to 800 nm in a fluorimeter (TECAN Spark 20M multimode microplate reader) with an excitation wavelength of 580 nm.

### PBS biotinylation

The ChromaLink™ Biotin Labeling reagent, purchased from Solulink (see https://vectorlabs.com/chromalink-biotin-antibody-labeling-kit.html), was used as it incorporates a chromophore as part of the linker. 120 μl of the PBS in 0.75 M potassium-phosphate pH 7.5 was buffer exchanged with the 1X Modification buffer (10X Modification buffer-100 mM sodium phosphate and 150 mM sodium chloride pH 8.0-diluted in 0.75 M potassium-phosphate pH 8.0). Then, the sample amount was quantified using an absorbance spectrum and using the values at 280, 354 and 620 nm to calculate the amount of biotin to add to the reaction. The samples were then incubated for 120 min at room temperature. The sample was buffer exchanged back into 100 μl of 0.75 M potassium-phosphate pH 7.5 using a Zeba Spin Desalt 7 kDa MWCO column, Thermo Scientific). UV–vis spectra were measured from 230–800 nm for biotinylated protein samples using a Tecan Safire microplate reader. Using the absorbance values at 280, 354 and 620 nm, and the molar extinction coefficients, the average number of biotin molecules covalently attached per PBS were calculated to be about 1. Fluorescence emission spectra were used to confirm functional energy transfer in the isolated PBS before and after biotinylation (Extended Data Fig. 1d).

### Cryo-EM sample preparation from purified PBS

For sample preparation we used Quantifoil Au 300 mesh 2/1 grids covered with a home-made streptavidin monolayer, which were manufactured as described previously^30^. To apply the sample, grids were first rehydrated in buffer A (375 mM potassium phosphate, pH 7.5) and then blotted dry with filter paper. 4 μl of PBS sample at 5.3 mg/ml in buffer B (750 mM potassium phosphate, pH 7.5) were added to the grid and then incubated on the bench for 60 seconds. Grids were washed on two 10 μl drops of buffer C (375 mM potassium phosphate, pH 7.5, 3% w/v trehalose, 0.01% v/v NP40, 0.05% w/v beta-octylglucoside) before being carefully wicked with Whatman filter paper. 1 μl of buffer C was added quickly, and the grid lifted into a FEI Mark IV Vitrobot. The grid was then manually blotted for 2-3s at 18°C and 100% humidity before plunging into a liquid ethane-propane (3:1) mix^44^. During the process, the PBS was exposed to ambient light for a total of approximately 2-3 min between pipetting and vitrification.

### Cryo-EM data collection PBS

For initial assessment of the sample, grids were loaded onto a Talos Arctica microscope (Thermo Fisher Scientific) equipped with a Gatan K3 direct electron detector and operating at 200 kV. 1954 movies were collected at a super-resolution pixel size of 0.558 Å, a defocus range of -0.6 μm to -1.8 μm and a total exposure of 50 e^-^/Å^2^ using SerialEM^45^. Images were processed as described below.

After evaluation of the first dataset, a second dataset was collected on a Titan Krios G3i microscope operating at an accceleration voltage of 300 kV, equipped with a Gatan K3 direct electron detector operating in CDS mode^46^ and a GIF Quantum energy filter with a 20-eV slit width. 12,051 movies were acquired with SerialEM in super-resolution counting mode with a super-resolution pixel size of 0.525 Å using an image shift collection scheme with active beam tilt correction, a defocus range from -0.5 μm to -1.6 μm and a total electron exposure of 50 e^-^/Å^2^.

### Image processing

All movies were aligned, gain corrected and binned by 2 using MotionCorr2 as implemented in RELION3^47^. The background streptavidin lattice in the motion corrected micrographs was subtracted using in-house scripts^30^. These subtracted micrographs were then imported into Cryosparc^48^ for patch CTF estimation and further image processing. For dataset 1 from the Talos Arctica microscope, particles were picked using the blob picker, extracted with a box size of 720 pixels, and subjected to one round of 2D classification. Good classes representing different views of the PBS were selected and used as templates for a second round of autopicking, yielding ∼156,000 particles. After another round of 2D classification, ∼43,000 particles in good classes were used to generate an ab-initio model of the PBS. A subsequent round of heterogenous refinement using this reference yielded two distinct classes representing the PBS^up-down^ (66 %) and PBS^up-up^ (34 %) conformations. These classes were refined to 4.1 Å and 4.5 Å, respectively.

For dataset 2 from the Titan Krios microscope, particles were picked from 800 micrographs using templates generated from the PBS^up-up^ reconstructions from dataset 1. After 2D classification and heterogenous refinement three distinct classes corresponding to PBS^up-down^, PBS^up-up^ and PBS^down-down^ conformations became apparent. Using these classes as references, 11,903 micrographs were picked to yield ∼1,900,000 particles. After several rounds of heterogenous refinement, approximately 510,000 good particles were sorted into four classes representing the three conformations (PBS^up-down^ 50.6%, PBS^up-up^ 38.9%, and PBS^down-down^ 10.5%). Refinement of the three conformations yielded overall resolutions of 2.7 Å (PBS^up-up^), 2.8 Å (PBS^up-down^) and 3.5 Å (PBS^down-down^).

During initial rounds of 2D classification of the datasets, we observed that several classes corresponded to PBS-rods that aligned independently of the core section of the PBS. To obtain reconstructions of the rods, these classes were used as templates for autopicking on a subset of 500 micrographs of dataset 2 and extracted with a box size of 360 pixels. 2D classification followed by ab-initio model generation and heterogenous refinement yielded a particle set that reached a resolution of 2.9 Å during non-uniform refinement. Using the same templates for picking from the entire dataset 2, ∼2,100,000 good particles were obtained that refined to a resolution 2.1 Å after CTF refinement, reaching Nyquist frequency. Further classification of the particle set to distinguish between different rod protomers within one PBS complex was unsuccessful and always yielding the same conformation. To visualize the contact area between two neighboring rods in the PBS^up-down^ conformation, particle subtraction followed by local refinement was carried out on the PBS^up-down^ reconstruction. The map yielded an improved resolution of 2.6 Å.

All processing was carried out using C1 symmetry and resolutions were estimated with the FSC = 0.143 criterion^49^. Reconstructions were sharpened using DeepEMhancer ^50^ with the default tight mask preset.

### Atomic model building and refinement

Using our PBS-OCP^R^ atomic models as a basis^41^, atomic models of the PBS core were refined using a cropped PBS^up-down^ map with the real space refinement program in PHENIX 1.19.2 (ref. ^51^). Similarly, models of the core of other PBS conformations were refined but found to be virtually identical. Models of the PBS-OCP^R^ rod were used as a basis for refinement of the PBS rod. To arrive at models for all three holo-PBS complexes, maps and models for the individual rods were rigid-body docked into the density of each holo-PBS conformer, which resulted in an unambiguous orientation of each rod. Because the holo-PBS maps are not resolved enough in the distal rod regions to allow for model refinement, models of the full complexes are for visualization only.

### LC-MS/MS analysis of the PBS SDS-PAGE fractions

Gel bands were digested in-gel according to^52^ with modifications. Briefly, gel bands were dehydrated using 100 % acetonitrile and incubated with 10 mM dithiothreitol in 100 mM ammonium bicarbonate, pH∼8, at 56 °C for 45 min, dehydrated again and incubated in the dark with 50 mM iodoacetamide in 100 mM ammonium bicarbonate for 20 min. Gel bands were then washed with ammonium bicarbonate and dehydrated again. Sequencing grade modified trypsin was prepared to 0.01 μg/μl in 50 mM ammonium bicarbonate and ∼100 μl of this was added to each gel band so that the gel was completely submerged. Bands were then incubated at 37 °C overnight. Peptides were extracted from the gel by water bath sonication in a solution of 60% Acetonitrile (ACN) /1% Trifluoroacetic acid (TFA) and vacuum dried to ∼2 μl.

Dried samples were re-suspended to 20 μl in 2% ACN/0.1% TFA and an injection of 5 μl was automatically made using a Thermo (www.thermo.com) EASYnLC 1000 onto a Thermo Acclaim PepMap RSLC 0.1mmx20mm C18 trapping column and washed for ∼5 min with buffer A. Bound peptides were then eluted over 35 min onto a Thermo Acclaim PepMap RSLC 0.075mm x 250mm resolving column with a gradient of 5%B to 40%B in 24 min, ramping to 90%B at 25 min and held at 90%B for the duration of the run (Buffer A = 99.9% water/0.1% formic acid, Buffer B = 80% acetonitrile/0.1% formic acid/19.9% water) at a constant flow rate of 300 nl/min. Column temperature was maintained at a constant temperature of 50 °C using and integrated column oven (PRSO-V1, Sonation GmbH, Biberach, Germany).

Eluted peptides were sprayed into a ThermoScientific Q-Exactive mass spectrometer (www.thermo.com) using a FlexSpray spray ion source. Survey scans were taken in the Orbitrap (70,000 resolution, determined at m/z 200) and the top 15 ions in each survey scan are then subjected to automatic higher energy collision induced dissociation (HCD) with fragment spectra acquired at 17,500 resolution. The resulting MS/MS spectra are converted to peak lists using Mascot Distiller, v2.7.1 (www.matrixscience.com), and searched against a database containing all *Synechocystis* PCC 6803 protein sequences and appended with common laboratory contaminants (downloaded 2020-11-17 from www.uniprot.org and www.thegpm.org, respectively) using the Mascot searching algorithm, v2.7. The Mascot output was then analyzed using Scaffold, v5.0 (www.proteomesoftware.com) to probabilistically validate protein identifications. Assignments validated using the Scaffold 1%FDR confidence filter are considered true.

Thee performed LC-MS/MS on the protein bands to confirm the identity of all the PBS components (Extended Data Table 2).

### Modelling the energy flow within PBS complex

Excitation energy transfer within PBS was studied in the limit of the Förster theory, using dipole-dipole approximation for the inter-pigment coupling. The energy transfer rates *k* (in ps^-1^) were computed using the equation^53^:

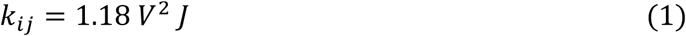

where *J* is the spectral overlap between donor emission and acceptor absorption, and the coupling *V* (in cm^-1^) is given by

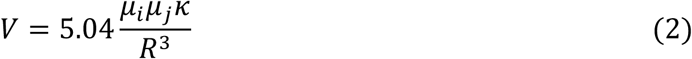

where *R* is the center-to-center dipole separation in nm, *k* is the orientation factor, and *μ_i_* and *μ_j_* are the transition dipole moments (in Debye) of the donor and acceptor, respectively. We have used effective transition dipole moments, including the correction for the effect of dielectric properties of the protein (see ref.^53^ for discussion of this point). The direction of the pigment transition dipoles of bilin chromophores were placed along the axis of the conjugated parts of the molecules. The geometrical axis was obtained using the singular-value decomposition on the atom coordinates. For bilins, the transition dipole moment of 13 D were used for the bilins in rod (CPC), and 15 D for bilins in the core (APC) pigments^54^. The spectral overlap *J* is calculated from the area-normalized spectra of donor and acceptor reported in literature^34, 55, 56^. Using the full matrix of the excitation transfer rates computed using (1), it is possible to write down a master equation describing the time evolution of probability of exciton residence on *i*-th chromophore, with a vector-valued *y*(*t*) = *yi*(t) (for *i* running from 1 to the number of pigments):

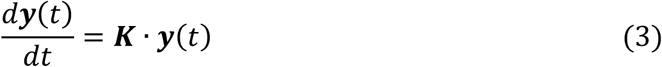

where ***K*** represents the transition matrix whose off-diagonal elements (*K_ij_*) represent the pairwise rates of energy transfer, *k*ij, between donor (*i*) and acceptor (*j*) pigments. The diagonal elements are defined as:

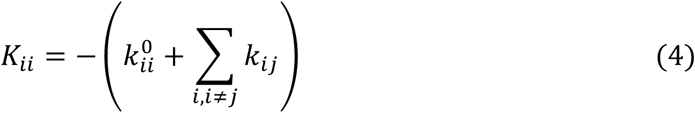

where 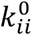 corresponds to the intrinsic excited state decay rate of the *i*-th pigment; we used 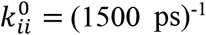 for all bilins, except for b-84, which was assigned the lifetime of 900 ps, based on^57^, however, the exact value of this parameter has little effect on the simulation results. Solving (3) for initial conditions given as *y_i_*(0) = 1 for *i* = *k*, and *y_i_*(0) = 0 for *i* ≠ *k*, yields the model for the excitation flow through the PBS following excitation into *k*-th pigment. A sum of *y_i_*(*t*) over *i* then describes the total decay of the excitation in the PBS. The validity of the model was tested by comparing the results with experimental data reported earlier. Extended Data Fig. 7 shows that correspondence between the simulation and experimental data is reasonable, considering that the model was not in any way adjusted to fit the experimental data and is based solely on the structural information and previously reported pigment spectra.

### Bioinformatics

Homologues of ApcG were found by searching with a profile HMM (constructed from sll1873 homologs obtained by BLAST search) against 420 non-redundant UniProt proteomes that contained a complete PBS protein set (as described in ref.^41^). Sequences of the 50 proteomes with a protein composition closest to *Synechocystis* PCC 6803 were used for sequence alignments. Sequences were aligned ClustalW^58, 59^, trimmed with trimAl^60^ and the protein sequence conservation was visualized with Weblogo 3.74^61^.Homologues of ApcG were found by searching with a profile HMM (constructed from sll1873 homologs obtained by BLAST search) against 420 non-redundant UniProt proteomes that contained a complete PBS protein set (as described in ref.^41^). Sequences of the 50 proteomes with a protein composition closest to *Synechocystis* PCC 6803 were used for sequence alignments. Sequences were aligned ClustalW^58, 59^, trimmed with trimAl^60^ and the protein sequence conservation was visualized with Weblogo 3.74^61^.

Structures were analyzed with ChimeraX^62^ and Pymol (The PyMOL Molecular Graphics System, version 1.7 Schrödinger, LLC).

## Acknowledgements

CAK and MADM would like to dedicate this manuscript to Dr. Nicole Tandeau de Marsac. The authors want to thank Dr. Bryan Ferlez for taking TEM images. We thank Dr. Douglas Whitten from the Proteomic Facility at Michigan State University, A. Chintangal and P. Tobias for computational support, Dr. Daniel Toso and Jonathan Remis at the Cal-Cryo facility for support with cryo-EM data collection, Dr. Robert Glaeser and Dr. Bong-Gyoon Han for advice concerning streptavidin grid preparation, and Dr. Lisa Eshun-Wilson for help with data processing. Research in the Kerfeld lab was supported by the Office of Science of the U.S. Department of Energy DE-FG02-91ER20021. This project has received funding from the European Union’s Horizon 2020 research and innovation programme under the Marie Sklodowska-Curie grant agreement No. 795070. DB and TP thank the Czech Science Foundation, grant No. 19-28323X. DB also acknowledges institutional support RVO:60077344. Molecular graphics and analyses were performed with UCSF ChimeraX with support from NIH R01-GM129325. E.N. is a Howard Hughes Medical Investigator.

## Author contributions

MADM and PVS designed and performed experiments, interpreted results. CK designed and supervised the project. HK helped with the sample preparation, interpreted results. MS refined the structures and interpreted results. DB and TP performed the model for the energy transfer. BJG performed initial characterization of the specimen for cryo-EM. MADM, PVS and CK wrote the manuscript with help from all authors.

## Competing interest

The authors declare no competing interests.

## Data availability

The atomic coordinates have been deposited in the Protein Data Bank with the accession codes 7SC8 for the rod and 7SC7 for the PBS^up-down^ core. The EM maps have been deposited in the Electron Microscopy Data Bank with the accession codes 25029 for the rod, 25028 for the PBS^up-down^ core, 25069 for the full map of the PBS^up-down^ conformation and the rod-rod contact, 25070 for the full map of the PBS^down-down^ conformation and 25071 for the full map of the PBS^up-up^ conformation.

All other data are available from the corresponding authors upon reasonable request.

## Extended Data

**Extended Data Table 1:** Synechocystis PCC 6803 Phycobilisome Protein subunits. (*) Nomenclature used along the text. (**) New linker discovered and incorporated in this study.

| Protein<br>component | Gene<br>symbol | Protein<br>name (*) | Function | pfam | #bilins per<br>monomer | #proteins<br>per PBS | MW (kDa) |
| --- | --- | --- | --- | --- | --- | --- | --- |
| α-PC | *cpcA* | CpcA | Rod<br>antenna<br>protein | pfam00502 | 1 PCB | 108 | 17.6 |
| β-PC | *cpcB* | CpcB | Rod<br>antenna<br>protein | pfam00502 | 2 PCB | 108 | 18.1 |
| L<sub>R33</sub> | *cpcC1* | CpcC1 | Internal rod<br>linker | pfam00427/<br>pfam01383 | -- | 6 | 32.5 |
| L<sub>R30</sub> | *cpcC2* | CpcC2 | Internal rod<br>linker | pfam00427/<br>pfam01383 | -- | 6 | 30.8 |
| L<sub>R10</sub> | *cpcD* | CpcD | Internal rod<br>linker | pfam01383 | -- | 6 | 9.3 |
| L<sub>RC</sub> | *cpcG1* | CpcG1 | Rod-to-core<br>linker | pfam00427 | -- | 6 | 28.9 |
| L<sub>RC</sub> | *cpcG2* | CpcG2 | Rod-to-core<br>linker | pfam00427 | -- | -- | 28.5 |
| α-APC | *apcA* | ApcA | Major core<br>antenna<br>protein | pfam00502 | 1 PCB | 32 | 17.4 |
| β-APC | *apcB* | ApcB | Major core<br>antenna<br>protein | pfam00502 | 1 PCB | 34 | 17.2 |
| L<sub>C</sub> | *apcC* | ApcC | Internal<br>core protein | pfam01383 | -- | 6 | 7.7 |
| L<sub>CM</sub> | *apcE* | ApcE | Core-to-<br>membrane<br>linker | pfam00427/<br>pfam00502 | 1 PCB | 2 | 100.3 |
| α-B | *apcD* | ApcD | Minor core<br>antenna<br>protein | pfam00502 | 1 PCB | 2 | 17.9 |
| α-B18 | *apcF* | ApcF | Minor core<br>antenna<br>protein | pfam00502 | 1 PCB | 2 | 18.9 |
| L<sub>C10</sub>** | *apcG* | ApcG | Linker | Newly<br>discovered<br>no assigned<br>pfam | -- | 2 | 12.9 |

**Extended Data Table 2:** MS analysis of the Synechocystis PCC 6803 Phycobilisome Protein subunits. The criteria were 2 peptides, False Discovery Rate (FDR) = 1%, percentage of coverage (% coverage) > 30%, and the number of total peptide count of each protein (# total peptide count). MW (kDa) = Molecular Weight assigned by the software used in the analysis, Scaffold 5.0 software.

| Protein<br>component | Gene symbol | # total peptide<br>count | % coverage | MW (kDa) |
| --- | --- | --- | --- | --- |
| α-PC | *cpcA* | 420 | 94 | 18 |
| β-PC | *cpcB* | 414 | 93 | 18 |
| L<sub>R33</sub> | *cpcC1* | 236 | 61 | 33 |
| L<sub>R30</sub> | *cpcC2* | 219 | 60 | 31 |
| L<sub>R10</sub> | *cpcD* | 148 | 88 | 9 |
| L<sub>RC</sub> | *cpcG1* | 215 | 83 | 27 |
| L<sub>RC</sub> | *cpcG2* | 21 | 43 | 29 |
| α-APC | *apcA* | 224 | 93 | 17 |
| β-APC | *apcB* | 322 | 94 | 17 |
| L<sub>C</sub> | *apcC* | 86 | 63 | 8 |
| L<sub>CM</sub> | *apcE* | 671 | 72 | 100 |
| α-B | *apcD* | 61 | 83 | 18 |
| α-B18 | *apcF* | 66 | 71 | 19 |
| L<sub>c10</sub> | *apcG* | 27 | 78 | 13 |
| FNR | *petH* | 177 | 78 | 46 |
| α-PC | *cpcA* | 420 | 94 | 18 |
| β-PC | *cpcB* | 414 | 93 | 18 |
| L<sub>R33</sub> | *cpcC1* | 236 | 61 | 33 |
| L<sub>R30</sub> | *cpcC2* | 219 | 60 | 31 |
| L<sub>R10</sub> | *cpcD* | 148 | 88 | 9 |
| L<sub>RC</sub> | *cpcG1* | 215 | 83 | 27 |
| L<sub>RC</sub> | *cpcG2* | 21 | 43 | 29 |
| FNR | *petH* | 177 | 78 | 46 |

**Extended Data Table 3:**
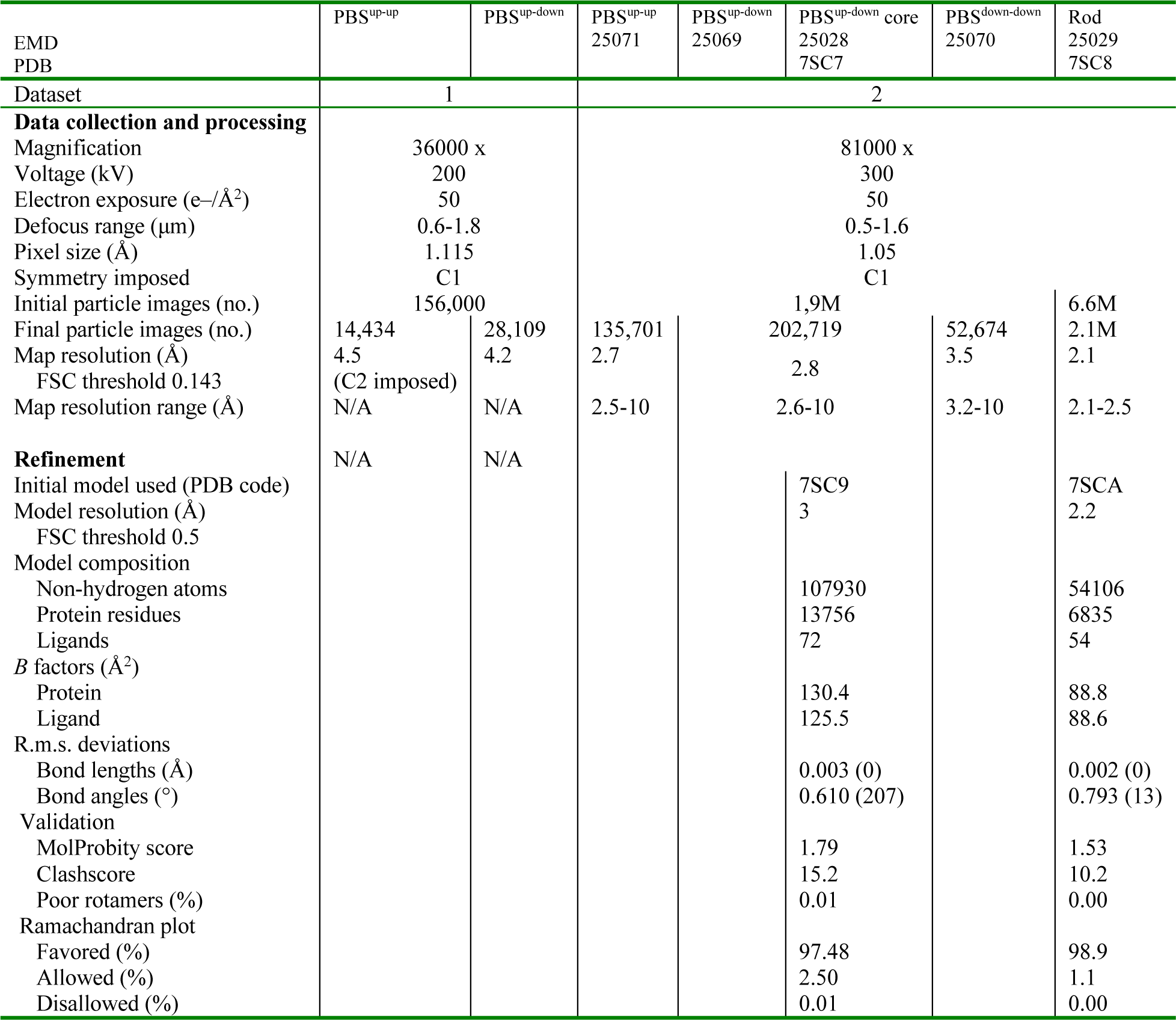
Cryo-EM data collection, refinement and validation statistics.

**Extended Data Figure 1:**
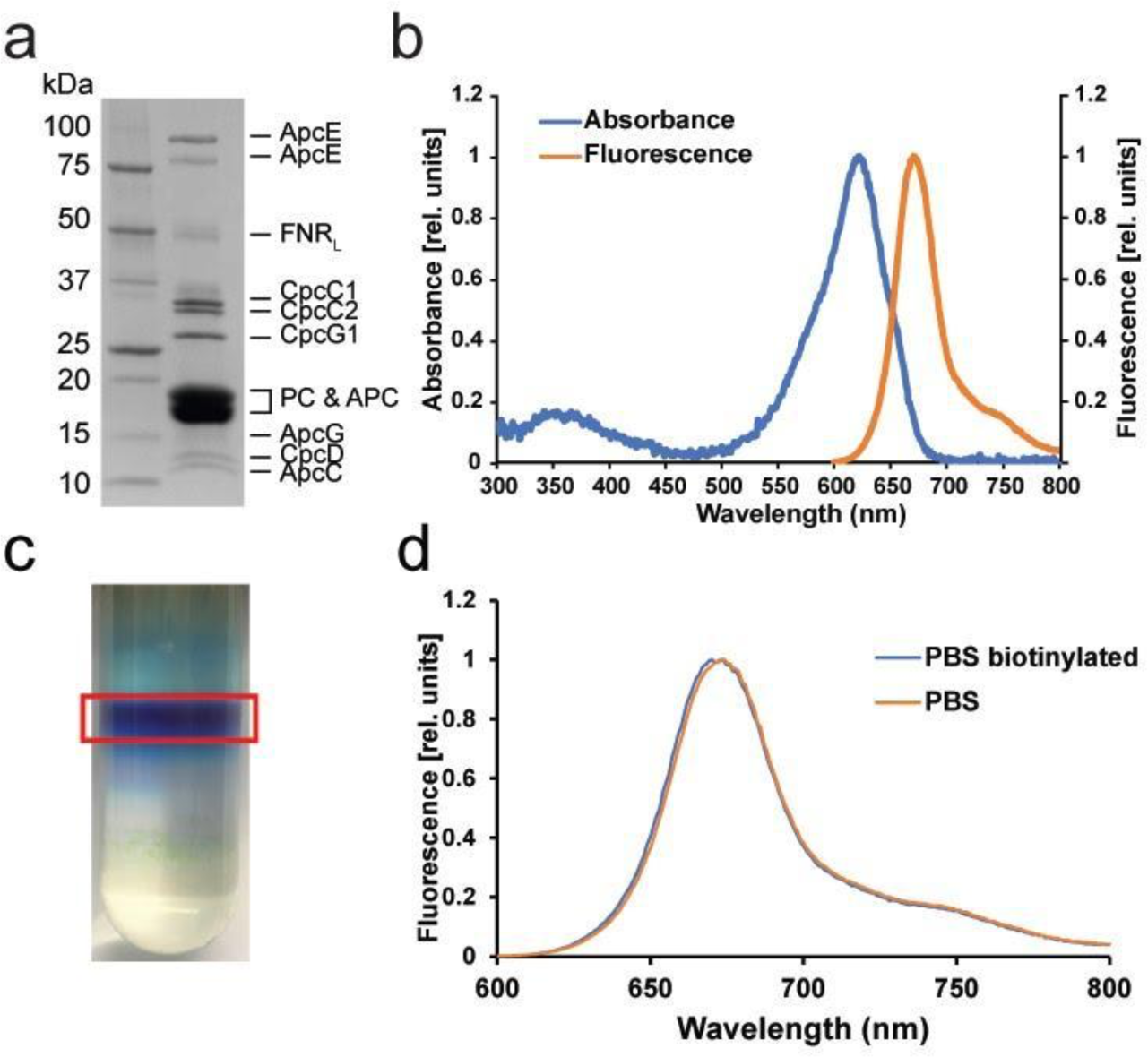
Biochemical and spectroscopic PBS characterization. **a**, Protein composition analysis of the PBS sample by SDS-PAGE. Identification of the components by MS can be found in Extended Data *Table 2*. **b**, Absorption, and fluorescence spectrum of the isolated PBS in 0.75 M K-phosphate pH 7.5. The PBS shows a maximum absorbance at 620 nm corresponding to phycocyanin (PC), and a shoulder at 650 nm corresponding to allophycocyanin (APC). The fluorescence peak maximum is ∼670 nm when PC is preferentially excited (excitation at 580 nm), and a shoulder from 740 nm to 780 nm. **c**, Sucrose gradient of PBS showing the intense blue band that was used for cryo-EM. **d**, Fluorescence spectra of PBS before and after in vitro biotinylation. Collectively, these data show that the PBS preparation was structurally and functionally intact.

**Extended Data Figure 2:**
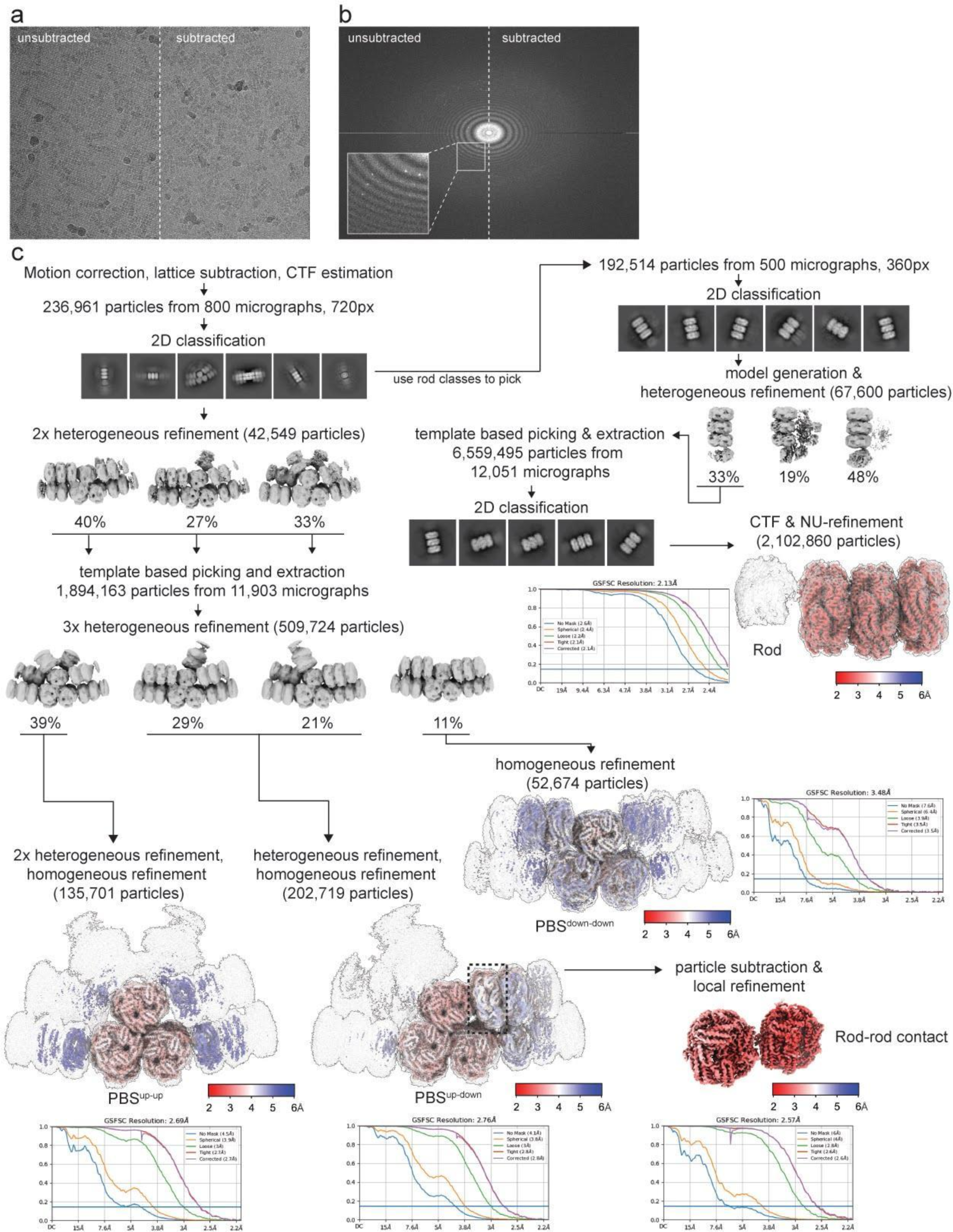
Cryo-EM data processing of PBS data. **a**, Example raw micrograph before and after streptavidin lattice subtraction. **b**, Fourier transform of **a** showing Bragg diffraction of streptavidin crystal before subtraction. **c**, Processing pipeline of dataset 2, yielding the structures presented in this paper. Red to blue color shadings represent local resolution estimates at FSC=0.5 ranging from 2 to 6 Å. The three PBS conformations and the structure of the rod are shown at two different thresholds to visualize both overall shape and higher resolution details.

**Extended Data Figure 3:**
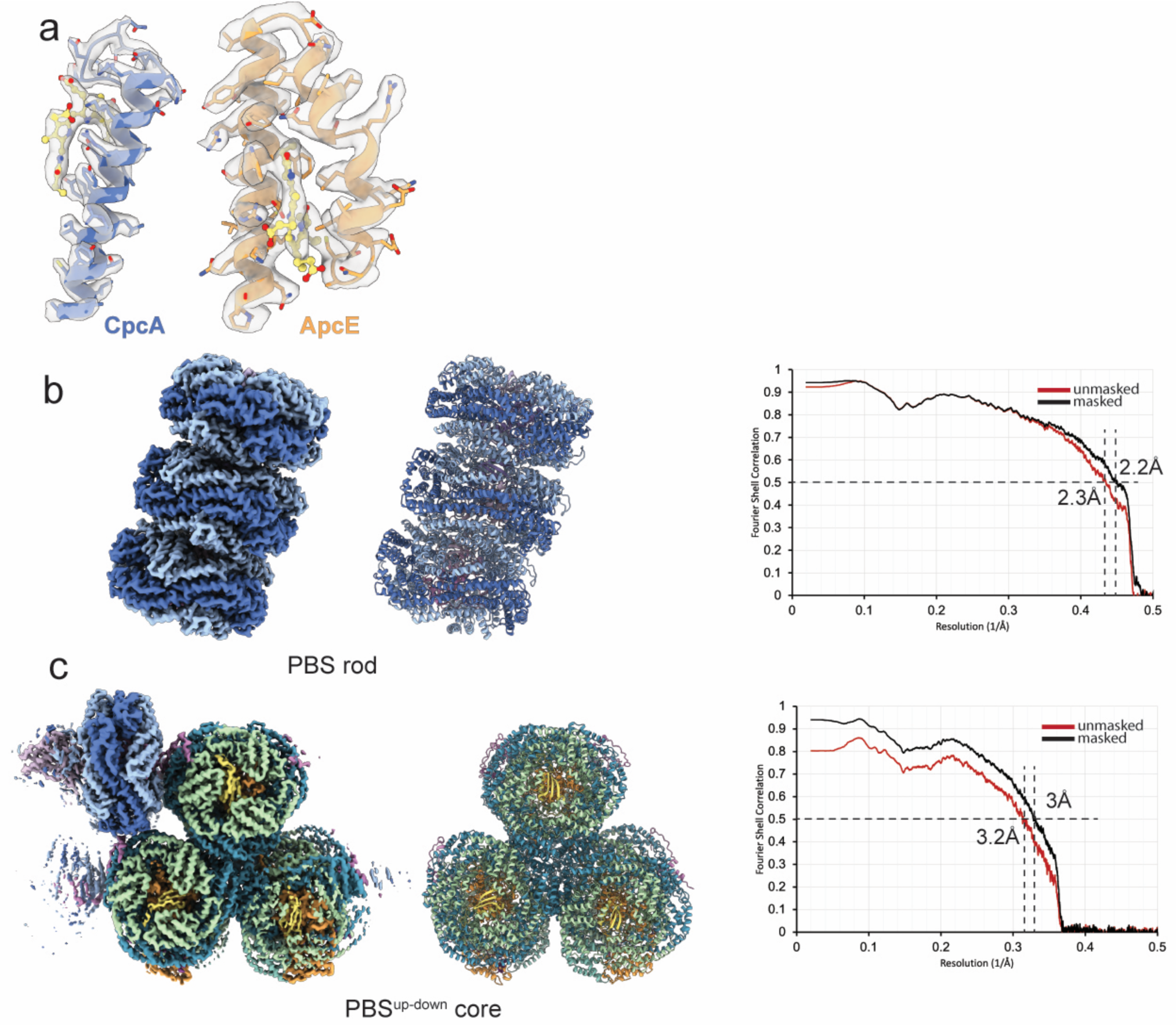
Map and Model quality of PBS^up-down^. **a**, EM density examples of CpcA, ApcE and associated bilins showing map quality of the PBS^up-down^. **b** and **c**, Cropped PBS^up-down^ map, model and corresponding Fourier shell correlation (FSC) curves. Resolution of masked and unmasked models are indicated at FSC = 0.5.

**Extended Data Figure 4:**
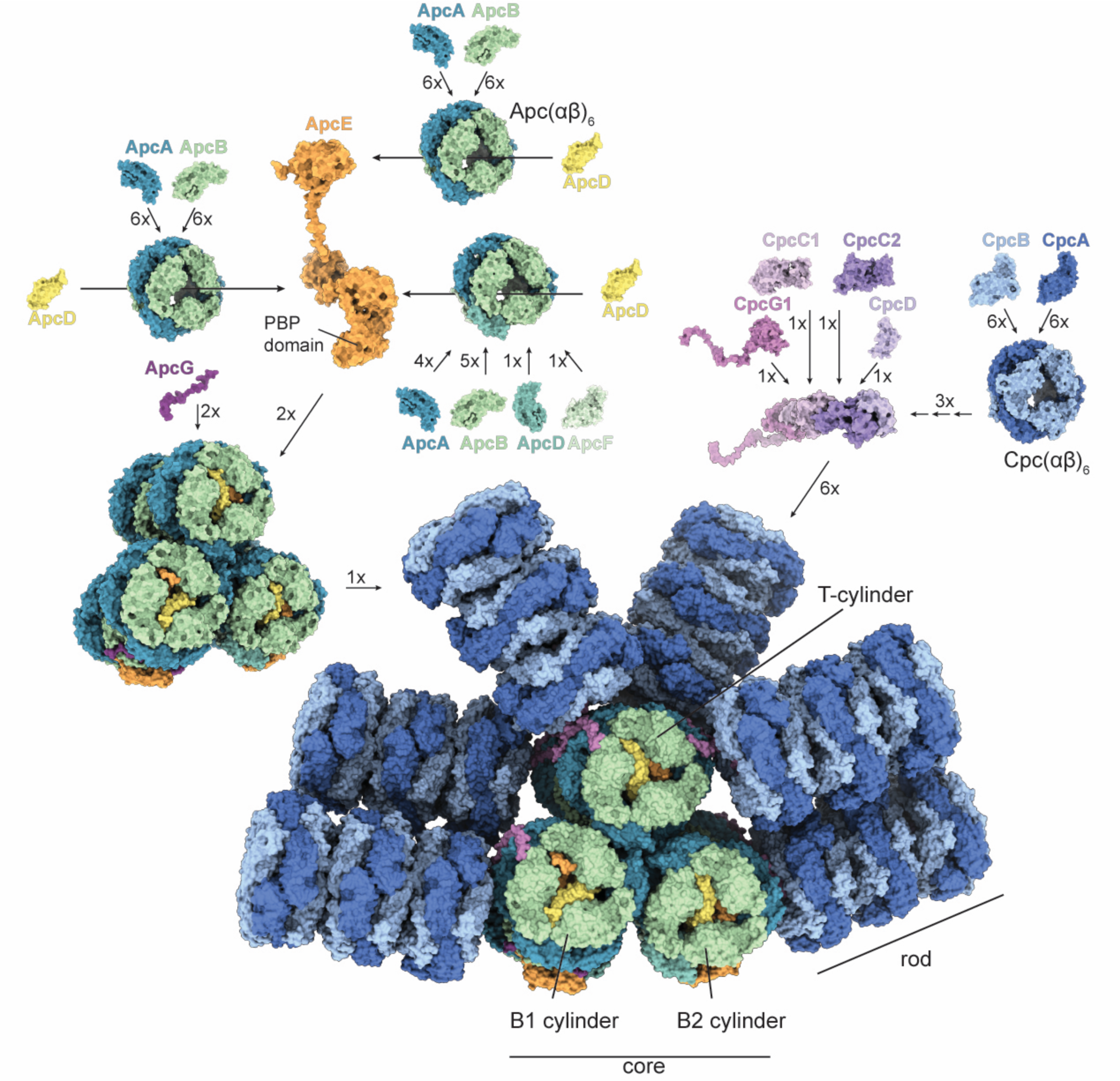
Assembly of the PBS. The core cylinders consist of four stacked discs; each disc contains three α- and three β-type Apc proteins that form the signature (αβ)6 hexamers. Besides the canonical ApcA and ApcB proteins, B1 and B2 cylinders contain one copy each of ApcD and ApcF, as well as the phycobiliprotein domain of ApcE. These are symmetrically arranged in the two bottom cylinders, oriented towards the membrane. ApcG. ApcD and ApcE protrude out of the core, indicating their likely role in contacting the photosystems. Each rod contains three stacked discs, each consisting of one CpcA/B hexamer. The overall architecture of the PBS core is in agreement with modelling predictions^29^.

**Extended Data Figure 5:**
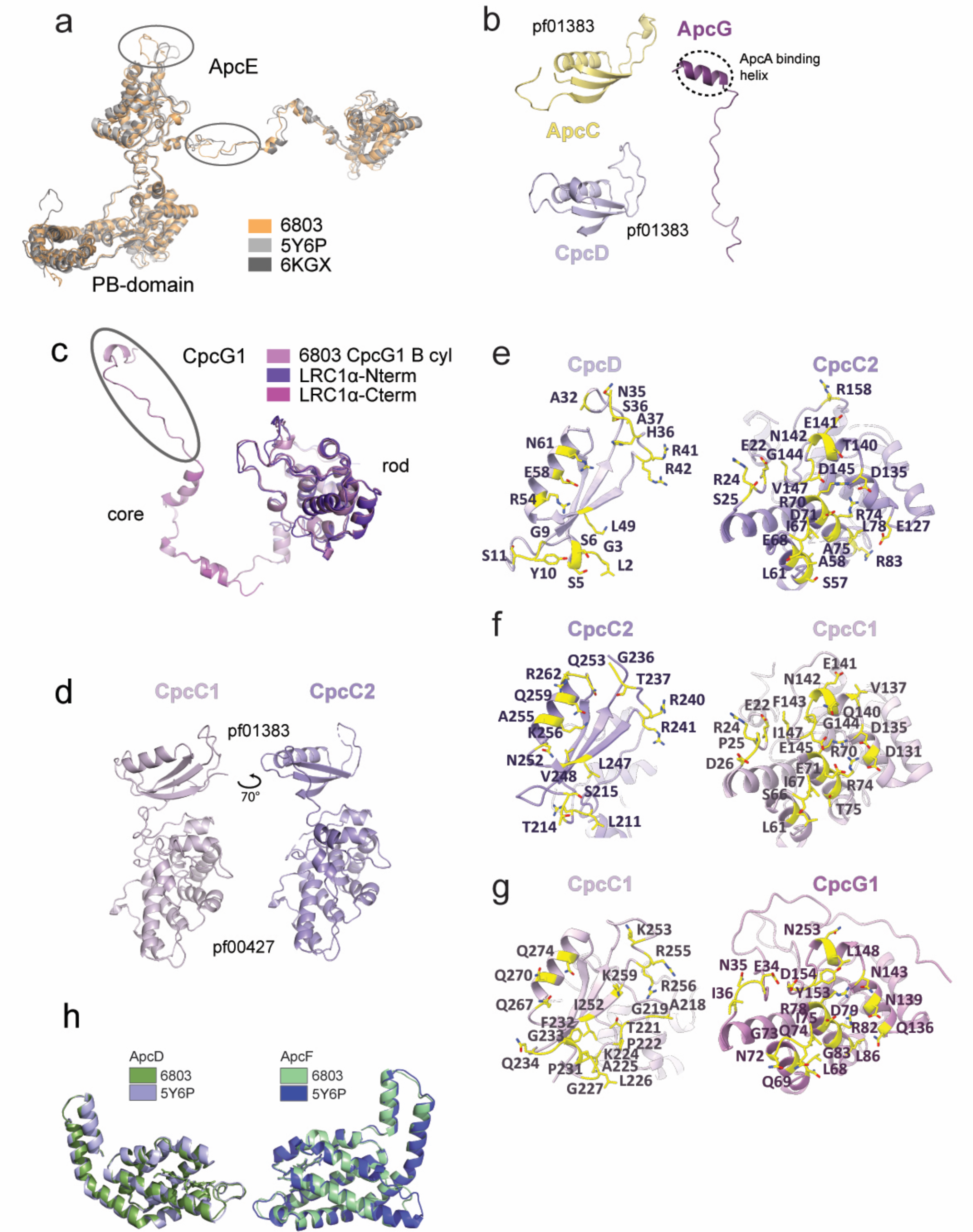
Molecular details of the PBS. **a**, Alignment of ApcE with red algae homologs (PDB:5Y6P Griffithsia pacifica, PDB:6KGX Porphyridium purpureum). RMSD is 1.9 Å, differences are circled. **b**, Structural models of ApcC, ApcG and CpcD. CpcD and ApcC consist of a single pf01383 domain. **c**, Alignment of CpcG1 with red algae homologs. Circle in CpcG1 shows C-terminal domain. CpcG1 is related to algal LRC1 linker with RMSD of 0.7 Å in the pf00427 domain. The C-terminal domain of CpcG1 has homology to LRC1a with RMSD of 0.6 Å. **d**, Comparison of CpcC1 and CpcC2. **e**-**g**, interaction between CpcD, CpcC2, CpcC1 and CpcG1. Interacting residues are highlighted. **h**, Alignment of ApcD and ApcF with red algae homologs. ApcD is related to the red algal homolog with RMSD of 0.6 Å while ApcF aligns with an RMSD of 0.6 Å with its homolog. The only major differences exist in loop regions.

**Extended Data Figure 6:**
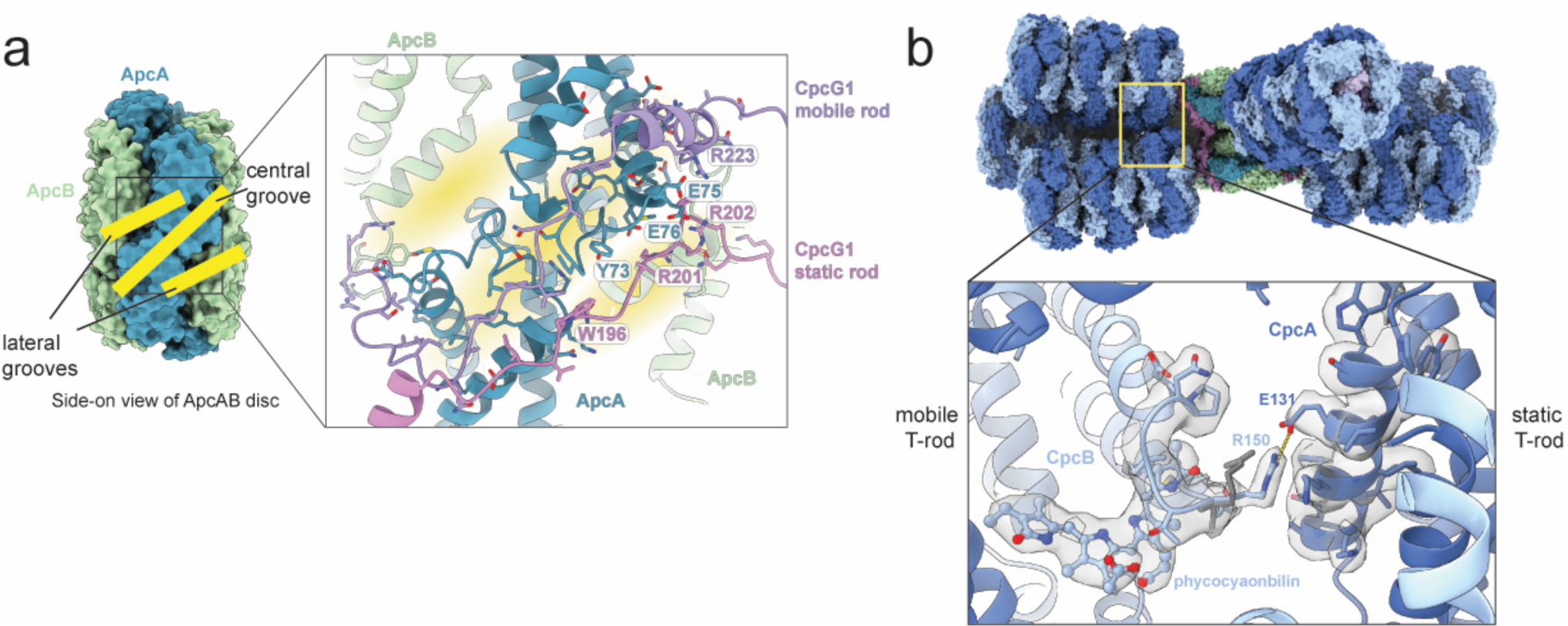
PBS rod conformation. **a**, Side view of AcpAB disc showing the groove network that organizes CpcG1 linker attachment to the PBS core. Selected residues are labeled. **b**, Salt bridge formed between two neighboring rods when the mobile rod adopts the ‘down’ conformation. EM density is transparent, dark grey and shows the conformation of R150 when the mobile rod is ‘up’. A bilin molecule in very close proximity of R150 of CpcB is also shown.

**Extended Data Figure 7:**
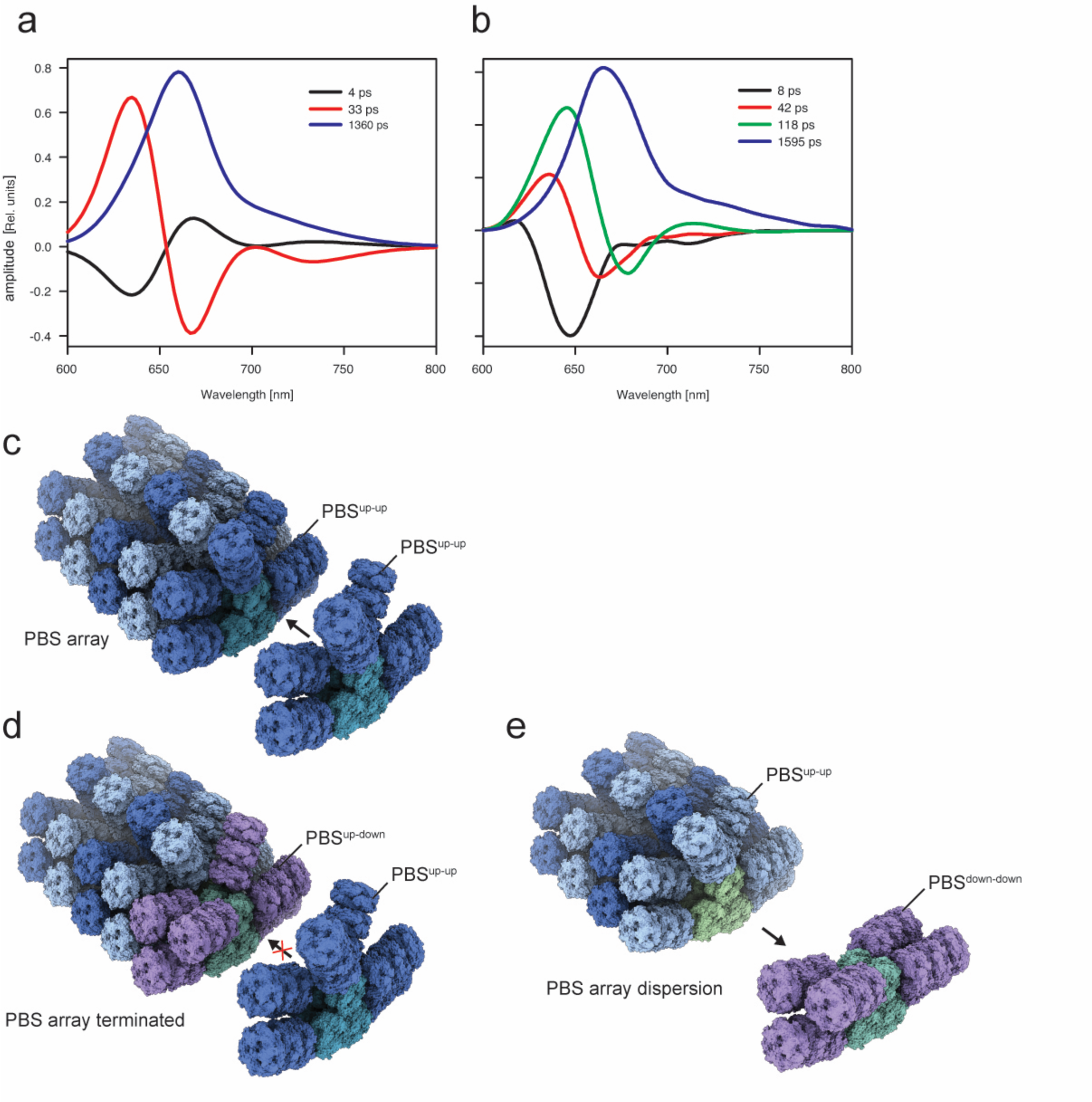
Decay associated spectra (DAS) and PBS arrays. DAS spectra obtained either **a**, from simulation or **b**, from fitting the data obtained from time-resolved fluorescence (right, adapted from ref.^34^) monitoring the excitation energy flow in the PBS. In simulations, PBS was excited into the far end of the rod. The ∼100 ps component (green) in the experimental data was interpreted as due to excitation annihilation^35^, hence it is not present in the simulation. **c**, PBS arrays were modeled based on cryo-electron tomography data from ref.^40^. a, only the up-up conformation is compatible with PBS arrays. **d**, The up-down conformation would result in array termination**. e**, A PBS switching to the down-down conformation could result in array dispersion.

## Notes

### Competing Interest Statement

The authors have declared no competing interest.

